# Ethological profiling of pain and analgesia in a mouse model of complex regional pain syndrome

**DOI:** 10.1101/2025.03.28.644648

**Authors:** Gabriella P. B. Muwanga, Amanda Pang, Sedona N. Ewbank, Janelle Siliezar-Doyle, Amy R. Nippert, Raag D. Airan, Vivianne L. Tawfik

**Author notes:** Correspondence should be addressed to: Vivianne L. Tawfik, Department of Anesthesiology, Perioperative & Pain Medicine, 3174 Porter Drive, Palo Alto, CA, 94304.

## Abstract

Complex regional pain syndrome (CRPS) is a form of chronic post-injury pain affecting the extremities. The mouse tibial fracture- cast model was developed to enable preclinical study of CRPS mechanisms and guide condition-specific drug development. Given the inherent limitations of reflex pain measures in mice, we sought to characterize pain-like behaviors in this model more holistically. We evaluated spontaneous and evoked pain and naturalistic behaviors after tibial fracture-cast injury in male mice in neutral and aversive environments using LabGym. Here, we report a unique ethological signature of pain in injured mice characterized by reflexive allodynia, thermal hyperalgesia, frequent grooming and reduced rearing in neutral and aversive environments, and decreased paw withdrawal and increased paw licking in an aversive environment. As proof-of-concept, we also leveraged this holistic behavioral evaluation for drug screening by characterizing the peripheral versus central effects of targeting alpha-2 receptors (α2-AR) in the tibial fracture-cast model. We evaluated the impact of systemic delivery of dexmedetomidine (DEX), a selective α2-AR agonist, with or without antagonists, on holistic behavioral metrics in injured male mice. We found that DEX reduced mechanical allodynia primarily via central α2-ARs. DEX also decreased motion metrics, grooming and rearing in an open field, and distinctly affected the quality and quantity of grooming in an aversive environment, and this effect was not suppressed by systemic α2-AR antagonists. Ultimately, this study holistically captures pain-related behaviors and provides a detailed characterization of the relative contributions of peripheral and central α2-ARs to alpha2-mediated analgesia in male mice after tibial fracture-cast injury.

## 1. INTRODUCTION

Complex regional pain syndrome (CRPS) is a form of chronic post-injury pain that develops in the extremities after minor trauma[56], affecting 50,000 new patients in the United States every year[58]. Understanding the pathophysiological mechanisms of CRPS is crucial for mechanism-based treatments. Tight casting of injured limbs and prolonged cast immobilization are established risk factors for CRPS development[13,54]. Based on this premise, the widely used mouse tibial fracture-cast model was developed to enable preclinical study of CRPS mechanisms[3] as it produces CRPS-like signs in mice that replicate key clinical features of the condition[67]. These include touch, heat and cold hypersensitivity, limb swelling and inflammation, osteopenia and loss of limb function, making this model a valuable tool for studying CRPS development and progression[3,16,28]. The model also serves as a platform to test potential therapies for this debilitating condition[3,23].

As with all pre-clinical pain models, reflex measures (ex. touch sensitivity/mechanical allodynia) are most reliably performed, but provide a limited evaluation of the “pain experience”, perhaps more specifically providing a readout of “nociception” after injury with limited translational validity[3,48,63]. Non-stimulus-evoked methods including grimace scales, locomotion, weight bearing and gait analysis can also be evaluated,[3,10,12,16,23,28,29,63] and specific to the CRPS model, peripheral signs such as changes in the color, temperature, and edema of the injured limb can be measured[3,16,29]. While these methods have proven useful in the preclinical setting, it is paramount that scientists also develop and utilize reliable and sensitive objective measures of spontaneous pain behavior to improve translatability of preclinical findings[48]. Although it has not hitherto been implemented in a CRPS model, pain may also be evaluated by manually scoring changes in naturalistic behaviors such as grooming and rearing, in which pain-induced changes occur in other models[12,14,65]. The advent of quantitative behavior analysis tools like DeepLabCut[46], DeepPoseKit[22], MoSeq[68], SLEAP[53], LabGym[27], SimBA[21], B-SoiD[26], VAME[42], etc., allow detailed and relatively unbiased evaluation of preclinical pain-induced spontaneous behaviors and naturalistic behaviors after injury[4]. LabGym[27], in particular, is user-friendly and allows for holistic assessment and quantification of diverse aspects of behavior.

Altered sweating, color, and skin temperature in the affected limb represent known sympathetic nervous system (SNS) involvement in CRPS[6]. Adrenergic receptors are the main effectors of SNS activation and the expression and efficacy of alpha-2 receptors (α2-AR), in particular, is altered in the peripheral and central nervous system after nerve injury[55]. Overall, the net effect of α2-AR activation is antinociception[45], making these receptors a viable analgesic target. The antinociceptive effects of α2-AR are predominantly attributed to the alpha 2a receptor subtype (α2A-AR) at supraspinal, spinal, or peripheral sites[44]. α2-AR are expressed in both sympathetic and primary afferent sensory neurons in the periphery. They inhibit exocytosis in these neurons by reducing calcium conductance into cells, preventing neurotransmitter release[72]. α2-ARs also have a major role in antinociception in the spinal cord dorsal horn[70]. In the spinal dorsal horn, the primary mechanisms for producing α2-AR-mediated pain suppression are presynaptic reduction of the release of excitatory acids from primary afferent central terminals[33,50] and the induction of inhibitory postsynaptic K+ current by direct action on α2-AR on spinal pain-relay neurons[62]. In physiologic conditions, the primary site of α2-AR agonist analgesic action is at the spinal cord[36], but this may change after injury. Limited clinical case series demonstrate that topical α2-AR agonists can decrease hyperalgesia, suggesting a peripheral effect[18,64], and central administration of α2-AR agonists reduces pain in patients with CRPS, suggesting a central effect[49]. However, the relative contributions of α2-AR signaling at each potential site of action in a sympathetically maintained pain model have not been explored.

The goals of this study were thus two-fold: 1) To holistically evaluate the tibial fracture-cast model using LabGym to assess changes in spontaneous and naturalistic behaviors in neutral and aversive environments, and 2) To evaluate the peripheral and central effects of α2-AR agonism on these behavioral metrics as proof-of-concept for therapeutic testing.

## 2. MATERIALS AND METHODS

### 2.1 Animals

Ten- to twelve-week-old adult male C57BL/6J (Jax stock #00664) mice, weighing approximately 25g, were used for all experiments. Each cage housed five mice in a room maintained on a 12-hour light/dark cycle in a temperature-controlled environment with ad libitum access to food and water.

### 2.2 Drugs and routes of administration

Dexmedetomidine hydrochloride (Sigma-Aldrich, #SML0956-10MG or #SML0956-50MG), Yohimbine hydrochloride (Abcam, # ab120239), Vatinoxan/MK-467 hydrochloride (MedChemExpress, #130466-38-5), and BRL-44408 maleate (Abcam, # ab120806) were dissolved in sterile water as directed by the manufacturer at a concentration of 10 mg/ml, 50 mg/ml, 5 mg/ml and 50 mg/ml respectively. Injured mice received a single dose of Dexmedetomidine hydrochloride intraperitoneally (IP, 1.25ug in 100ul; 50ug/kg). For antagonist experiments, injured mice were injected intraperitoneally with either yohimbine hydrochloride (0.05mg in 100ul; 2mg/kg), Vatinoxan/MK-467 hydrochloride (0.05mg in 100ul; 2mg/kg), or BRL-44408 maleate (0.05mg in 100ul; 2mg/kg) 30 minutes before injecting Dexmedetomidine hydrochloride (1.25ug in 100ul; 50ug/kg) intraperitoneally.

### 2.3 Tibial fracture-cast model of complex regional pain syndrome (CRPS)

Mice were anesthetized with isoflurane and underwent a closed right distal tibia fracture followed by casting. The right hind limb was wrapped in gauze and a hemostat was used to make a closed fracture of the distal tibia. The hind limb was then wrapped in casting tape (ScotchCast™ Plus) from the metatarsals of the hind paw up to a spica formed around the abdomen to ensure that the cast did not slip off. The cast over the paw was applied only to the plantar surface with a window left open over the dorsum of the paw and ankle to prevent constriction when post-fracture edema developed. Mice were inspected throughout the postoperative period of cast immobilization to ensure that the cast was properly positioned. At three weeks post-fracture, mice were briefly anesthetized, and casts were removed with cast shears. For behavioral assessment, mice were tested beginning one day after cast removal at three weeks until six days after cast removal. CRPS model generation and behavioral testing were conducted following well established methods for evaluating mouse behavior in the tibial fracture-cast model of CRPS[3,10,16,23,28,29].

### 2.4 Behavioral testing

All testing was conducted within the same time window on each testing day in an isolated, temperature- and light-controlled room. Mice were acclimated for 60 minutes in the testing environment within custom clear plastic cylinders (4” D) on a raised metal mesh platform (24” H). To ensure blinding of the experimenter, mice were either randomly placed in a cylinder and identified after testing, or mice were assigned to a group and injected with a coded treatment whose identity was revealed to the experimenter after the experiment was complete and the data were processed.

### 2.5 Mechanical nociception assays

To evaluate mechanical reflexive hypersensitivity, we used a logarithmically increasing set of 8 von Frey filaments (Stoelting), ranging in gram force from 0.007 to 6.0 g. These were applied perpendicular to the plantar hind paw with sufficient force to cause a slight bending of the filament. A positive response was characterized as a rapid withdrawal of the paw away from the stimulus filament within 4 s. Using the up-down statistical method, the 50% withdrawal mechanical threshold scores were calculated for each mouse and then averaged across the experimental groups.

### 2.6 Paw Edema, Unweighting and Temperature Measurements

Hind paw edema was determined by measuring the hind paw dorsal-ventral thickness over the midpoint of the third metatarsal with digital calipers while the mouse was restrained in a tube. Hind paw thickness data were analyzed as the difference between the fracture side and the contralateral intact side and averaged across experimental groups. Paw edema was measured at 3 weeks post-fracture one hour before drug injection and two hours after drug injection.

An incapacitance device (IITC Inc Life Science, Woodland Hills, CA) was used to measure hind paw unweighting. Mice were manually held in a vertical position over the apparatus with the hind paws resting on separate metal scale plates, and the entire weight of the mouse was supported on the hind paws. The duration of each measurement was 6 seconds, and 6 consecutive measurements were taken at 60-second intervals. Six readings were averaged to calculate the bilateral hind paw weight-bearing values. Unweighting was measured at baseline and then again at 3 weeks post-fracture, and 1- and 2-hours post DEX injection. Data were analyzed as a ratio between the right hind paw weight divided by the sum of right and left hind paw values [2R/(R + L)] × 100%).

The temperature of the hind paw was measured using a fine-gauge thermocouple wire (Omega, Stamford, CT). Temperature testing was performed over the hind paw dorsal skin between the first and second metatarsals (medial), the second and third metatarsals (central), and the fourth and fifth metatarsals (lateral). The measurements for each hind paw were averaged for the mean paw temperature. Data were expressed either as temperature in degrees Celsius (°C) or as the average difference between the ipsi- and contralateral hind paw within an experimental group. Paw temperature was measured at three weeks post-fracture before and after (1 and 2 hours) drug injection.

### 2.7 Thermal nociception assays

To evaluate thermal-induced reflexive responses, we used the Hargreaves and hotplate tests.

For the Hargreaves test, mice were placed in their glass bottomed enclosures and a radiant heat source aimed at the plantar surface of the mouse’s right hind paw was positioned beneath the animal. The time taken to withdraw from the heat stimulus was manually recorded as the paw withdrawal latency in seconds. A cut off time of 30 seconds was set to prevent thermal injury to the paw. Paw withdrawal latencies were recorded at baseline and three weeks post-fracture.

For the hot plate test, the plate was set to 50°C, mice were placed on the plate and recorded for 45 seconds using a high definition DFK 22AUC03 video camera (60 fps). Only one exposure to the hotplate was applied on a given testing session to prevent behavioral sensitization that can result from multiple noxious exposures. In cases where multiple measurements were done on the same cohort of mice, at least two days of rest were left between testing sessions. Measurements were done at baseline and 3 weeks post-fracture.

### 2.8 Open field video recordings and analysis

All open field recordings were taken overhead using a high definition DFK 22AUC03 video camera (30 fps) in two side-by-side 45cmx45cmx45cm boxes. Mice were recorded at baseline, post-fracture and one-hour post-drug injection for 30 minutes at each timepoint. Before any analysis was done, videos were cropped so that each video contained one box with one animal.

#### Mouse tracking and ROI analysis

Mice were tracked and ROI analysis was performed using AnyMaze (https://www.any-maze.com/). Briefly, videos were imported into AnyMaze, and two ROIs were defined and manually drawn on each video. The first ROI was a square measuring 25cmx25cm representing the center zone, and the second ROI was a larger square measuring 30cmx30cm demarcating the peripheral thigmotaxis zone. We then proceeded with ROI analysis and generated CSV files containing descriptive statistics for distance traveled (m), average speed, time spent immobile, number of entries into each ROI, and the time spent in each ROI.

#### Behavior stratification using LabGym

Open field videos were also analyzed using LabGym[27]. To analyze different behaviors of interest, models were trained as outlined on the LabGym GitHub page (https://github.com/umyelab/LabGym). A detailed behavior analysis pipeline is depicted in Figure 1. To analyze different behaviors of interest, example frames were generated using LabGym and uploaded to Roboflow (app.roboflow.com). 90 frames were annotated in Roboflow and used to train a detector for open field videos in LabGym. The trained open field mouse detector was used to generate behavior bout examples (15 frames per bout) which were manually sorted into the following behavior classes: body grooming, face grooming, walking, rearing and idling. Sorted examples were then used to train a categorizer whose performance was tested on other sets of independently sorted data as shown in Supplementary figure 1. The trained categorizer was used to generate behavioral categorizations with 40% uncertainty that were then fed into the quantifier to compute specific quantitative measurements for different aspects of each behavior as defined in Hu et al. (2023)[27].

**Figure 1.**
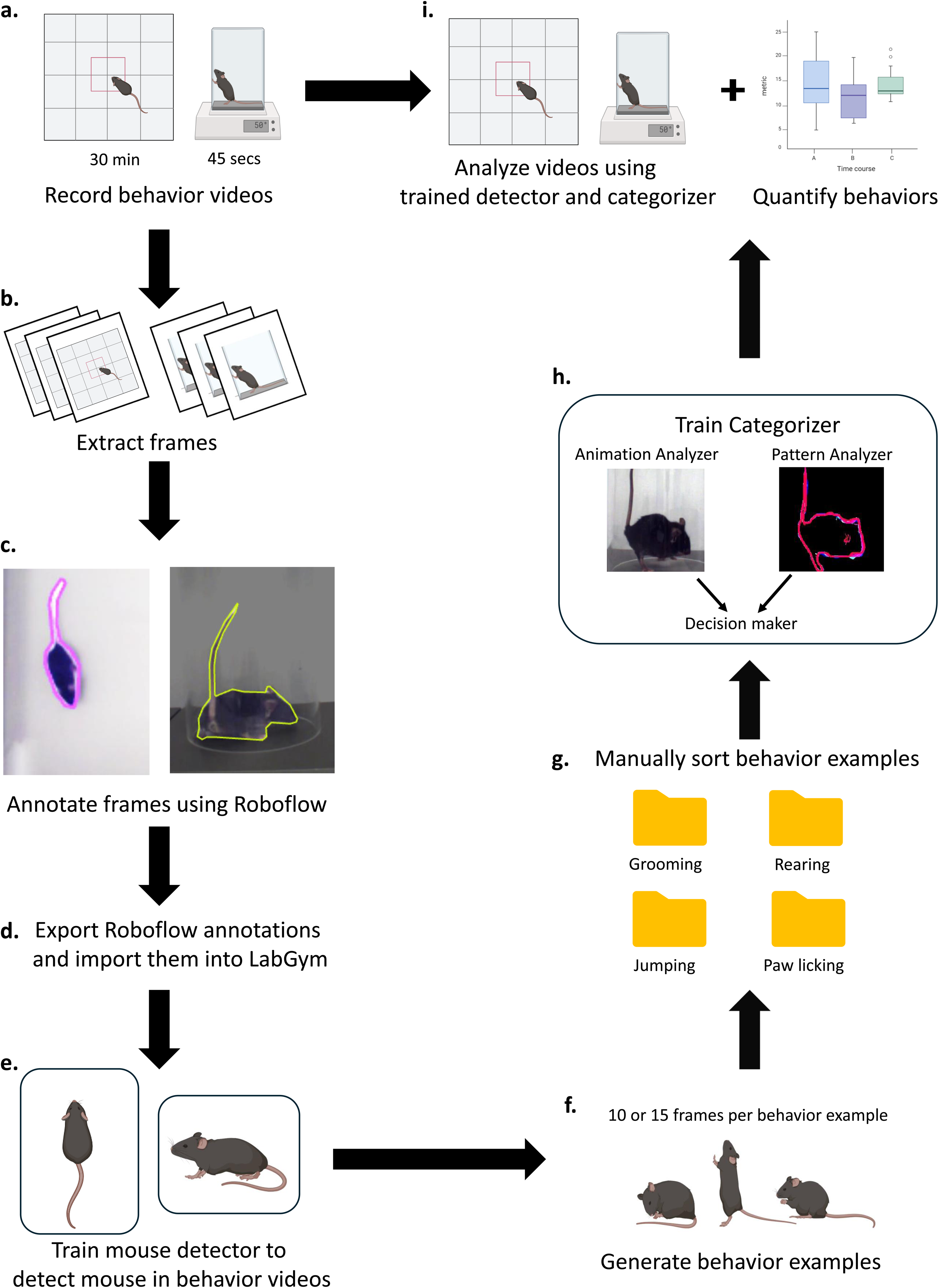
The behavior video analysis pipeline. **a.** Behavior videos are recorded. **b.** A representative sample of frames is extracted from recorded videos. **c.** Extracted frames are annotated using Roboflow. **d.** Annotated frames are extracted and imported into LabGym. **e.** Annotations are used to train a mouse detector in LabGym. **f.** The mouse detector is used to generate behavior examples from recorded videos (10 or 15 frames per behavior example). **g.** Behavior examples are manually sorted into behaviors of interest. **h.** Sorted behavior examples are used to train a behavior categorizer. **i.** Test videos are analyzed using the trained detector and categorizer and behaviors of interest are quantified.

Quantitative behavior measures generated in this study as defined by LabGym: Count: The frequency of a behavior for the entire duration of analysis[27].

Duration: The total time spent performing a specific behavior[27].

Latency: The time it takes for a behavior to occur for the first time during analysis[27].

Intensity: The summary of how intense a behavior is calculated as the proportional changes of the animal body area between frames during a behavior divided by the time window for categorizing the behaviors[27].

Magnitude: The summary of the magnitude of motion calculated as the maximum proportional change in animal body area when performing a specific behavior[27].

Vigor: The summary of how vigorous behavior is calculated as which is the magnitude divided by the time in which a behavior bout takes place[27].

### 2.9 Hotplate video analysis

As with open field video recordings, hotplate videos were analyzed using LabGym[27]. Briefly, example frames were generated using LabGym and uploaded to Roboflow (app.roboflow.com). A total of 914 frames were annotated to locate the mouse using instance segmentation in Roboflow. Annotations were exported from Roboflow, imported into LabGym and used to train a highly precise and accurate detector in LabGym. The trained detector was used to generate behavior bout examples (10 frames per bout) which were manually sorted into the following behavior classes: grooming, jumping, paw withdrawal, paw licking, rearing, and other (representing miscellaneous behaviors observed). 1540 sorted bout examples were then used to train a categorizer, whose performance was also evaluated on independently sorted data. The trained categorizer was used to generate behavioral categorizations with 40% uncertainty which were then quantified, exported and plotted using GraphPad Prism version 10.2.2.

### 2.10 Quantification and statistical analysis

Measurements of cohort sizes were determined based on historical data from our laboratory using a power analysis to provide >80% power to discover 25% differences with p<0.05 between groups to require a minimum of 4 animals per group for all behavioral outcomes. All experiments were randomized by cage and performed by a blinded researcher. Researchers remained blinded throughout behavioral assessments. Groups were unblinded at the end of each experiment after statistical analysis. All data are expressed as the mean ± SEM. Statistical analysis was performed using GraphPad Prism version 10.2.2 (GraphPad Software). Where appropriate, data were analyzed using ordinary one-way ANOVA with a Tukey correction, ordinary two-way ANOVA with a Tukey or Sidak correction, and two-tailed Student’s t tests. No data was excluded from analyses.

### 2.11 Study Approval

All procedures were approved by the Stanford University Administrative Panel on Laboratory Animal Care in accordance with American Veterinary Medical Association guidelines and the International Association for the Study of Pain.

## 3. RESULTS

### 3.1 The tibial fracture-cast model results in a CRPS-like syndrome in mice that reproduces clinical signs

Prior to deep ethological studies, we sought to confirm the development of common CRPS signs in our experimental mice by evaluating mechanical sensitivity, thermal sensitivity, weight bearing, paw thickness and temperature as shown in Figure 2a. As previously established in the lab[16,29], we found that tibial fracture-cast injury resulted in a significant increase in mechanical sensitivity indicated by a decreased paw withdrawal threshold to mechanical stimuli (Figure 2b). Tibial fracture-cast injury also reduced the paw withdrawal latency to a focused thermal stimulus when tested three weeks post-fracture, indicating an increase in thermal sensitivity in injured mice (Figure 2c). Injured mice also placed less weight on the injured than the uninjured paw (Figure 2d). There was also a significant increase in injured paw width (Figure 2e) and temperature (Figure 2f) compared to the uninjured paw, indicating post-injury edema and inflammation.

**Figure 2.**
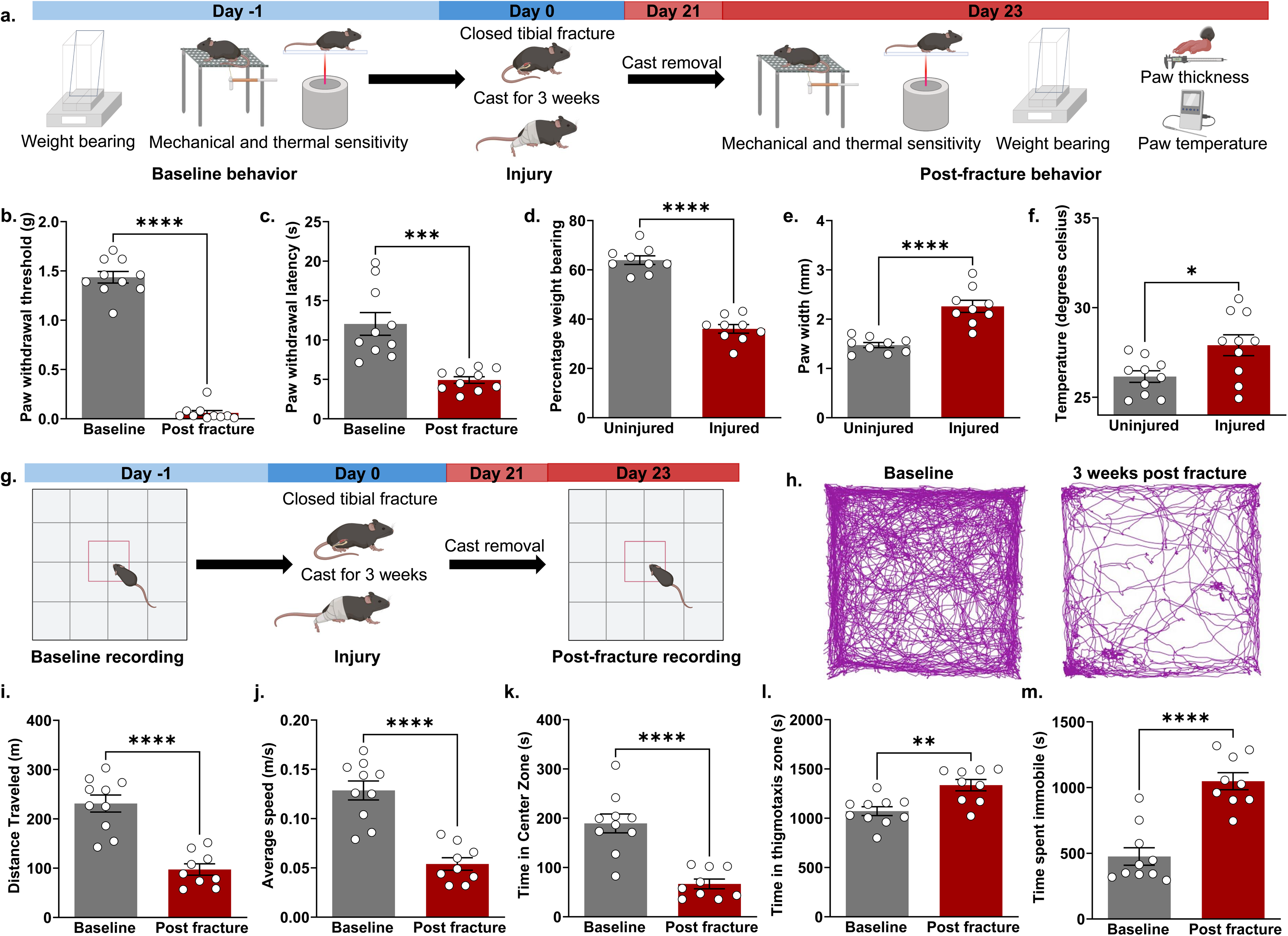
Tibial fracture-cast injury reproducibly results in CRPS-like symptoms and motion changes in mice. **a.** Schematic diagram showing experimental timeline for behavior testing (n= 10 male mice). Tibial fracture-cast injury increases mechanical (**b**) and thermal sensitivity (**c**) while decreasing weight bearing on the injured paw (**d**) and increasing paw edema (**e**) and temperature (**f**) at three weeks post-fracture. **g.** Schematic diagram showing experimental timeline for open field behavior testing (n= 10 male mice). **h.** Representative path plots showing post-injury changes in motion in injured mice. Tibial fracture-cast injury reduces distance traveled (**i**) and average speed (**j**) in an open field three weeks post-fracture. Tibial fracture-cast injury also reduces time spent in the center zone (**k**) while increasing time spent in the thigmotaxis zone (**l**) of the open field, and time spent immobile (**m**) three weeks post-fracture. All behavior data shown in this figure was analyzed using Student’s t-test (*p<0.05, **p<0.01, ***p<0.001, ****p<0.0001). Data are expressed as the mean ± SEM.

We next characterized the effect of tibial fracture-cast injury on motion in an open field to evaluate mobility/function as outlined in Figure 2g. We observed that CRPS mice traveled shorter distances (Figure 2h-i) at lower average speeds (Figure 2j) after injury compared to their pre-injury baseline. Injured mice also spent less time in the center zone (Figure 2k) and more time in the thigmotaxis zone (Figure 2l) of the arena. Generally, test mice also spent more total time immobile after injury (Figure 2m).

We therefore concluded that tibial fracture-cast injury results in a CRPS-like syndrome that recapitulates common signs including paw edema, increased paw temperature, mechanical allodynia, thermal hyperalgesia, anxiety-like behaviors, and impaired mobility.

### 3.2 Semi-supervised behavior analysis reveals distinct changes in naturalistic behaviors in a neutral environment after tibial fracture-cast injury

First, we evaluated our LabGym open field model on three manually sorted datasets namely an naïve/uninjured animal dataset, injured animal dataset, and a mixed (naïve/uninjured and injured) dataset. The LabGym open field model performed well on all datasets, achieving high precision, recall and F1 scores for all behaviors of interest as shown in Supplementary Figure 1.

Next, we conducted an in-depth analysis of specific naturalistic behaviors in an open field and the impact of tibial fracture-cast injury on those behaviors (Figure 3a). We observed an increase in the proportion of animals grooming their faces and bodies, and a decrease in the proportion of animals walking and rearing over the course of the recording (Figure 3b). Our analysis also revealed an increase in how much and how long injured mice groomed their faces compared to their pre-injury baseline (Figure 3c). Injury also significantly decreased face grooming intensity and vigor three weeks post fracture (Figure 3c). Similarly, tibial fracture-cast injury increased body grooming counts and duration, and decreased body grooming intensity and vigor (Figure 3d). Injured mice also walked less frequently and spent less time walking with a coinciding reduction in walking intensity and vigor (Figure 3e). Lastly, we found that injury decreased rearing bout count, bout duration, vigor and intensity (Figure 3f). However, tibial fracture-cast injury did not affect any behavior latencies (Supplementary figure 2a). These results show that tibial fracture-cast injury distinctly affects the quality and quantity of naturalistic behaviors in a neutral environment and that these changes can be reliably identified using robust learning-based holistic assessment.

**Figure 3.**
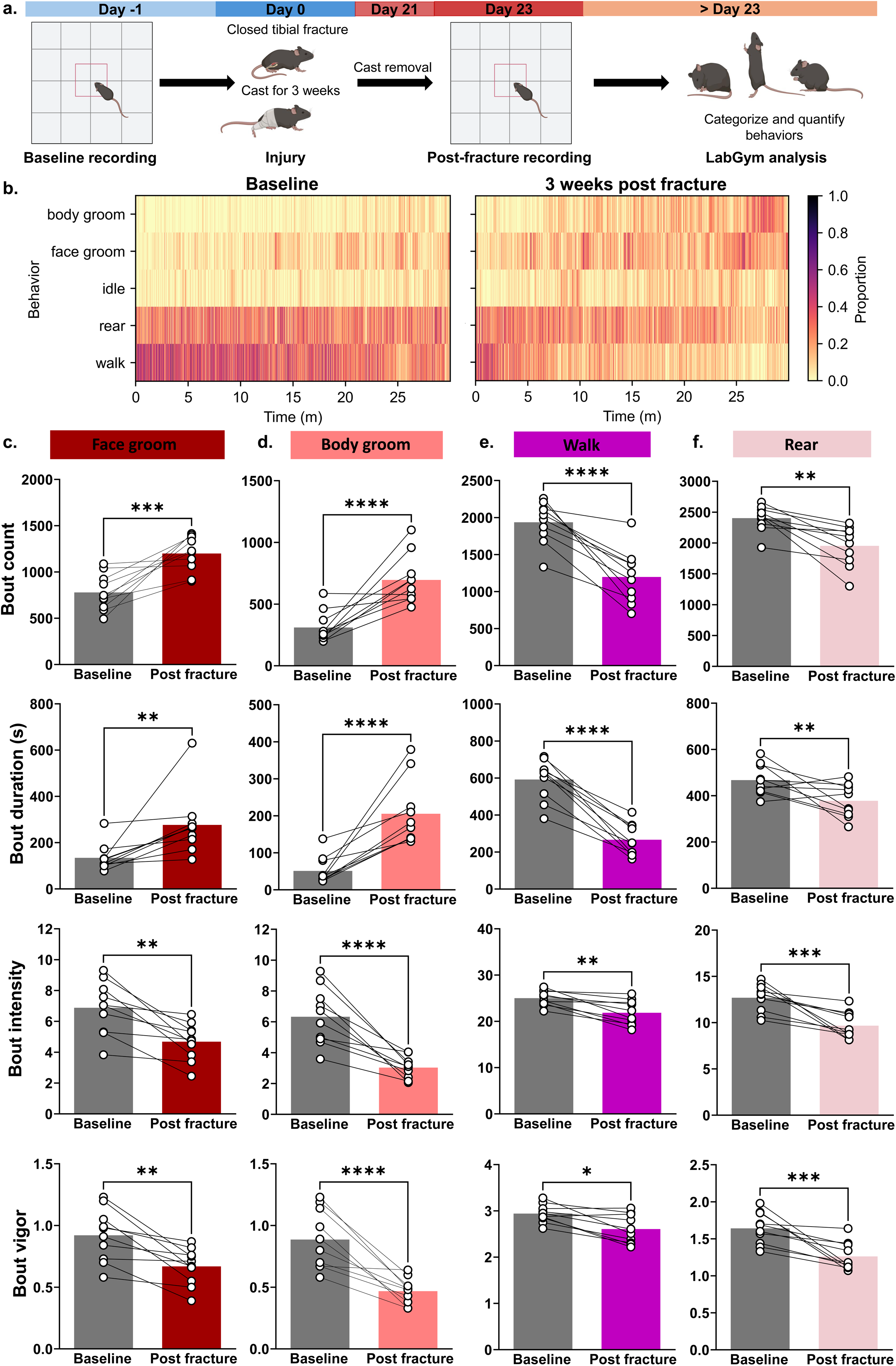
Tibial fracture-cast injury in mice causes distinct changes in naturalistic behaviors in an open field. **a.** Schematic diagram showing experimental timeline for open field behavior testing and analysis (n= 10 male mice). **b.** Representative raster plots showing proportions of mice performing each behavior over the course of the analysis. **c.** Tibial fracture-cast injury increases face grooming frequency and duration and reduces face grooming intensity and vigor. **d.** Tibial fracture-cast injury increases the frequency and duration of body grooming and decreases body grooming intensity and vigor. **e.** Tibial fracture-cast injury decreases walking bout count, duration, intensity and vigor. **f.** Tibial fracture-cast injury decreases rearing count, duration, intensity and vigor. All behavior data shown in this figure was analyzed using Student’s t-test (*p<0.05, **p<0.01, ***p<0.001, ***p<0.0001). Data are expressed as the mean ± SEM.

### 3.3 Semi-supervised behavior analysis reveals distinct changes in naturalistic and nocifensive behaviors in a noxious heat environment after tibial fracture-cast injury

As was done with naturalistic behaviors in an open field, we leveraged the power of machine learning to evaluate the effect of a noxious heat stimulus on naturalistic (rearing, grooming) and pain-related behaviors (paw withdrawal, paw licking, jumping) (Figure 4a). Our LabGym hotplate analysis model achieved high levels of accuracy for each of our behaviors of interest as shown in Supplementary figure 3. The animals in this cohort did not jump while on the hotplate for the duration of analysis so this behavior was excluded from further analysis.

**Figure 4.**
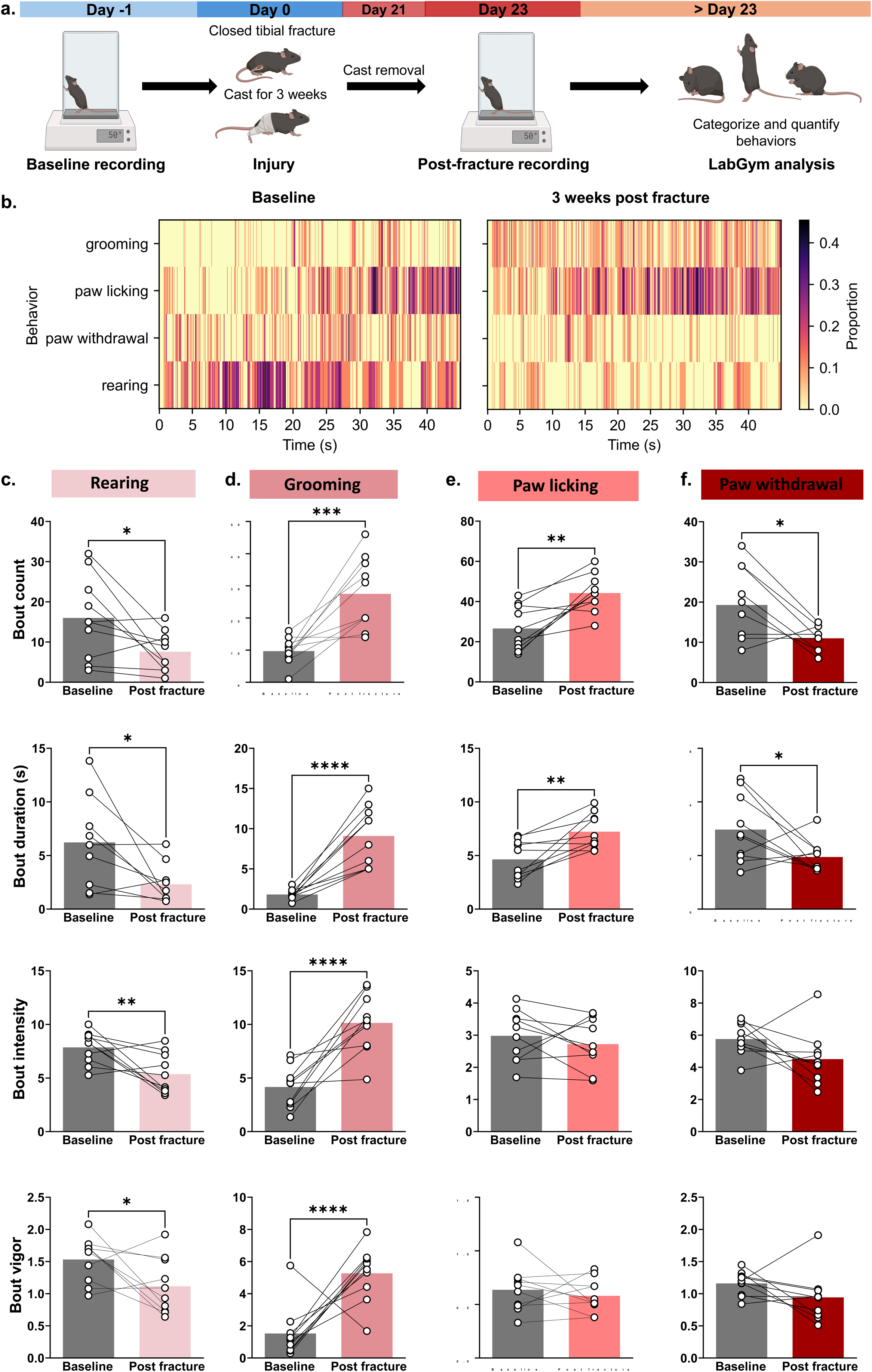
Tibial fracture-cast injury in mice causes distinct changes in pain-related behaviors in a noxious heat environment. **a.** Schematic diagram showing experimental timeline for behavior testing (n= 10 male mice). **b.** Representative raster plots showing proportions of mice performing each behavior over the course of the analysis. **c.** Tibial fracture-cast injury reduces rearing count, duration, intensity and vigor in a noxious heat environment. **d.** Tibial fracture-cast injury increases grooming count, duration, intensity and vigor in a noxious heat environment. **e.** Tibial fracture-cast injury increases paw licking count and duration but does not affect paw licking intensity and vigor in a noxious heat environment. **f.** Tibial fracture-cast injury reduces paw withdrawal count and duration but does not affect paw withdrawal intensity and vigor in a noxious heat environment. All behavior data shown in this figure were analyzed using Student’s t-test (*p<0.05, **p<0.01, ***p<0.001, ***p<0.0001). Data are expressed as the mean ± SEM.

We observed an increase in the proportion of animals grooming and licking their paws, with a concomitant decrease in the proportion of animals withdrawing their paws and rearing over the course of the recording (Figure 4b). Our analysis also revealed that tibial fracture-cast injury decreased rearing count, duration, intensity and vigor (Figure 4c), but did not affect rearing latency. Tibial fracture-cast injury also increased grooming counts, duration, intensity and vigor (Figure 4d) but grooming latency was unaffected. While paw licking count and duration were also higher after injury, paw licking intensity and vigor were lower after injury (Figure 4e). Paw licking latency also apparently increased but this effect was not statistically significant (P = 0.0551) (Supplementary figure 4a). We also discovered that injured mice withdrew their paws from the hotplate less frequently with a matching decrease in paw withdrawal duration (Figure 4f). However, paw withdrawal intensity, vigor and latency remained unchanged (Figure 4f, Supplementary figure 4a respectively). Miscellaneous behaviors (categorized as ‘other’) were mostly unchanged after injury, but their vigor and intensity reduced (Supplementary figure 4b). Collectively, these results show that tibial fracture-cast injury changes the nature of naturalistic and nocifensive behaviors in a noxious heat environment which can be reliably identified using robust machine learning-based holistic assessment.

### 3.4 α2-AR agonism alleviates pain after tibial fracture-cast injury

After successfully characterizing behavior changes after tibial fracture-cast injury, we sought to characterize the effect of targeting α2-ARs on evoked and spontaneous pain behaviors, motor function and anxiety-like behaviors in injured mice. We used dexmedetomidine (DEX), an imidazole compound that displays specific and selective α2-AR agonism, to target α2-ARs for the subsequent studies.

First, we investigated the effect of intraperitoneal DEX injection on mechanical sensitivity in fractured mice three weeks post-fracture (Figure 5a). We found an increase in mechanical withdrawal thresholds starting at 30 minutes and lasting up to 120 minutes after intraperitoneal DEX injection (Figure 5b), indicating reversal of mechanical allodynia. Next, we investigated the effect of α2-AR agonism on paw edema, weight bearing and paw temperature three weeks post fracture. We were intrigued to find that systemic DEX had no effect on post-fracture weight bearing (Figure 5c) and paw width in injured mice three weeks post-fracture (Figure 5d). It, however, significantly reduced the temperature of the injured paw 1-hour post-injection in CRPS mice three weeks post-fracture (Figure 5e).

**Figure 5.**
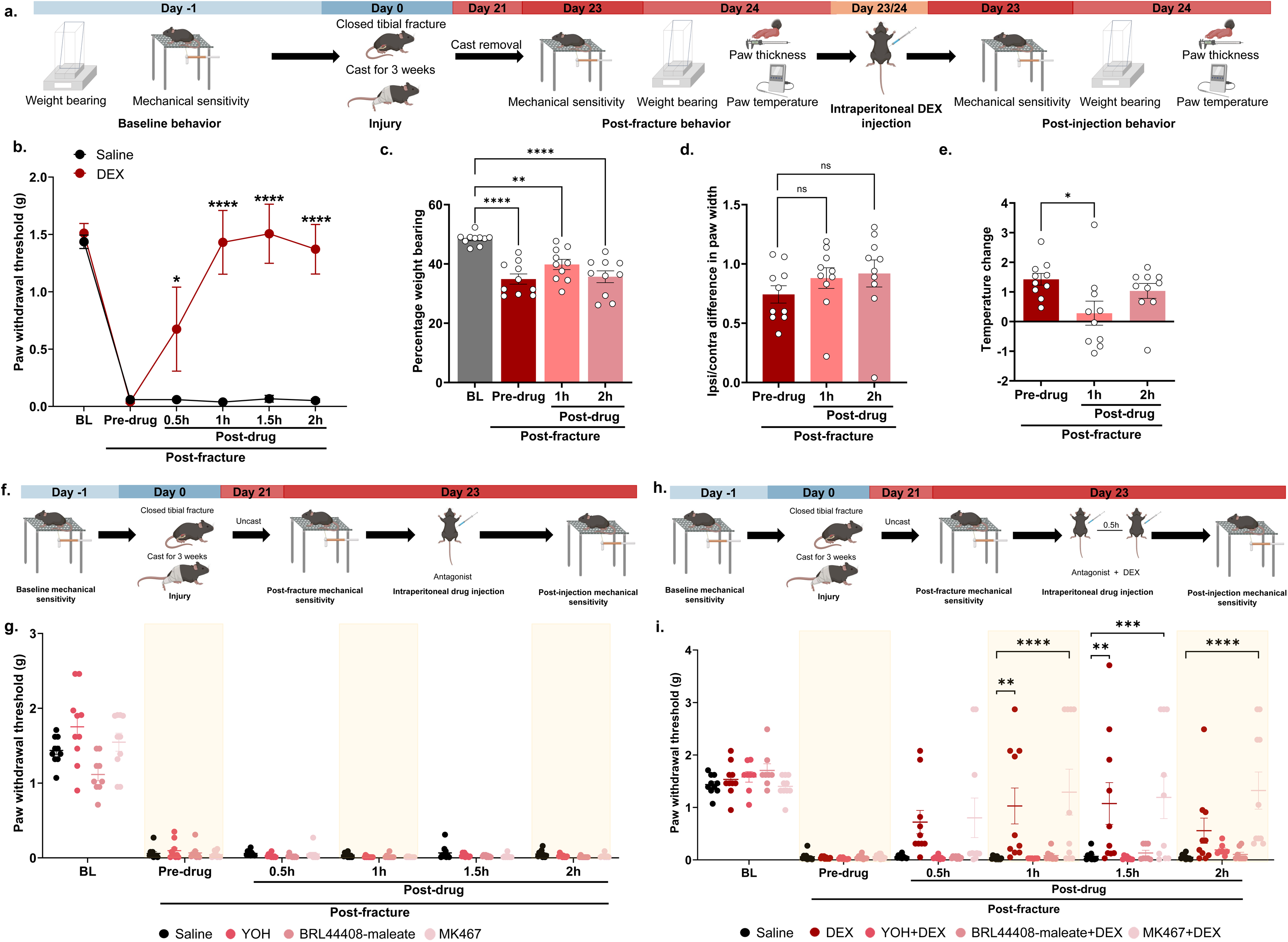
Systemic α2-AR agonism reduces reflex behaviors and paw warmth via central action. **a.** Schematic diagram showing experimental timeline for open field behavior testing and analysis. **b.** Intraperitoneal dexmedetomidine (DEX) reverses mechanical allodynia three weeks post-fracture (two treatment groups: saline or DEX; n= 10 male mice per group). Intraperitoneal DEX does not affect percentage weight bearing on the injured paw (**c**) and paw width (**d**), but reduces paw temperature (**e**) at three weeks post fracture (single treatment group: DEX; n= 10 male mice). **f.** Schematic diagram showing experimental timeline for the α2AR antagonist behavioral experiments (n= 10 male mice per group). **g.** Neither yohimbine (YOH), BRL44408 maleate, nor MK467 alone affect mechanical allodynia in injured mice. **h.** Schematic diagram showing experimental timeline for the α2AR antagonist + DEX behavioral experiments (n= 10 male mice per group). **i.** Administering DEX in the presence of YOH and BRL44408 maleate, but not MK467, reverses DEX-mediated effects on mechanical allodynia in injured mice. All behavior data in this figure was analyzed using an ordinary two-way ANOVA with Tukey correction (*p<0.05, **p<0.01, ***p<0.001, ****p<0.0001). Data are expressed as the mean ± SEM.

Next, we further interrogated the site of action and the relative contributions of central and peripheral α2-AR to the anti-allodynic effects seen in prior experiments. To ascertain that antagonists alone did not affect mechanical allodynia in injured mice, we administered either a systemic α2-AR antagonist (yohimbine hydrochloride), a systemic α2A-AR antagonist (BRL-44408 maleate) or a peripherally restricted α2A-AR antagonist (Vatinoxan/MK-467) (Figure 5f). None of the adrenergic antagonists used affected mechanical allodynia three weeks post-fracture when administered alone (Figure 5g). We then administered either yohimbine hydrochloride, BRL-44408 maleate or Vatinoxan/MK-467 thirty minutes before intraperitoneal DEX injection (Figure 5h). When administered with DEX, yohimbine hydrochloride and BRL-44408 maleate reversed DEX’s anti-allodynic effects (Figure 5i). Interestingly, Vatinoxan/MK-467 did not affect DEX-mediated improvement in mechanical allodynia (Figure 5i), suggesting that the analgesic site of action of systemic DEX might be in the central nervous system.

Having evaluated DEX effects on mechanical sensitivity and autonomic symptoms in mice three weeks after injury, we proceeded to investigate the effects of α2-AR agonism on motion dynamics in an open field. Administering DEX systemically after injury further reduced the distance traveled and average speed but did not significantly affect post-injury reduction in time spent in the center zone (Supplementary figure 5d, e, f respectively). DEX administration after injury also significantly increased thigmotaxis zone time and immobility time in injured mice (Supplementary figure 5g and h respectively).

Taken together, these results reveal centrally mediated α2-AR effects on mechanical allodynia after injury. They also depict mild sedation of DEX-treated mice that affects motion in an open field, possibly masking any analgesic effect.

### 3.5. α2-AR agonism alters grooming and rearing in an open field after tibial fracture-cast injury

The next step was to evaluate the effect of targeting α2-AR on naturalistic behaviors in an open field as shown in Figure 6a. We found that administering DEX to injured mice reduced the proportion of injured animals grooming, rearing, and walking over the course of analysis (Figure 6b). Systemic DEX also further reduced face grooming frequency, intensity and vigor (Figure 6c), but did not affect face grooming duration (Figure 6c). DEX injection also decreased body grooming counts and duration, but did not affect body grooming intensity and vigor after injury (Figure 6d). We also observed a reduction in walking bout counts and duration post DEX injection with no accompanying drug-induced change in walking intensity and vigor after injury (Figure 6e). Rearing bout frequency, duration, intensity and vigor also decreased more after DEX treatment in injured mice (Figure 6f). Generally, DEX treatment did not alter behavior latencies after injury (Supplementary figure 6a). Although mice apparently spent more time idle after DEX treatment, this change was not statistically significant (Supplementary figure 6b).

**Figure 6.**
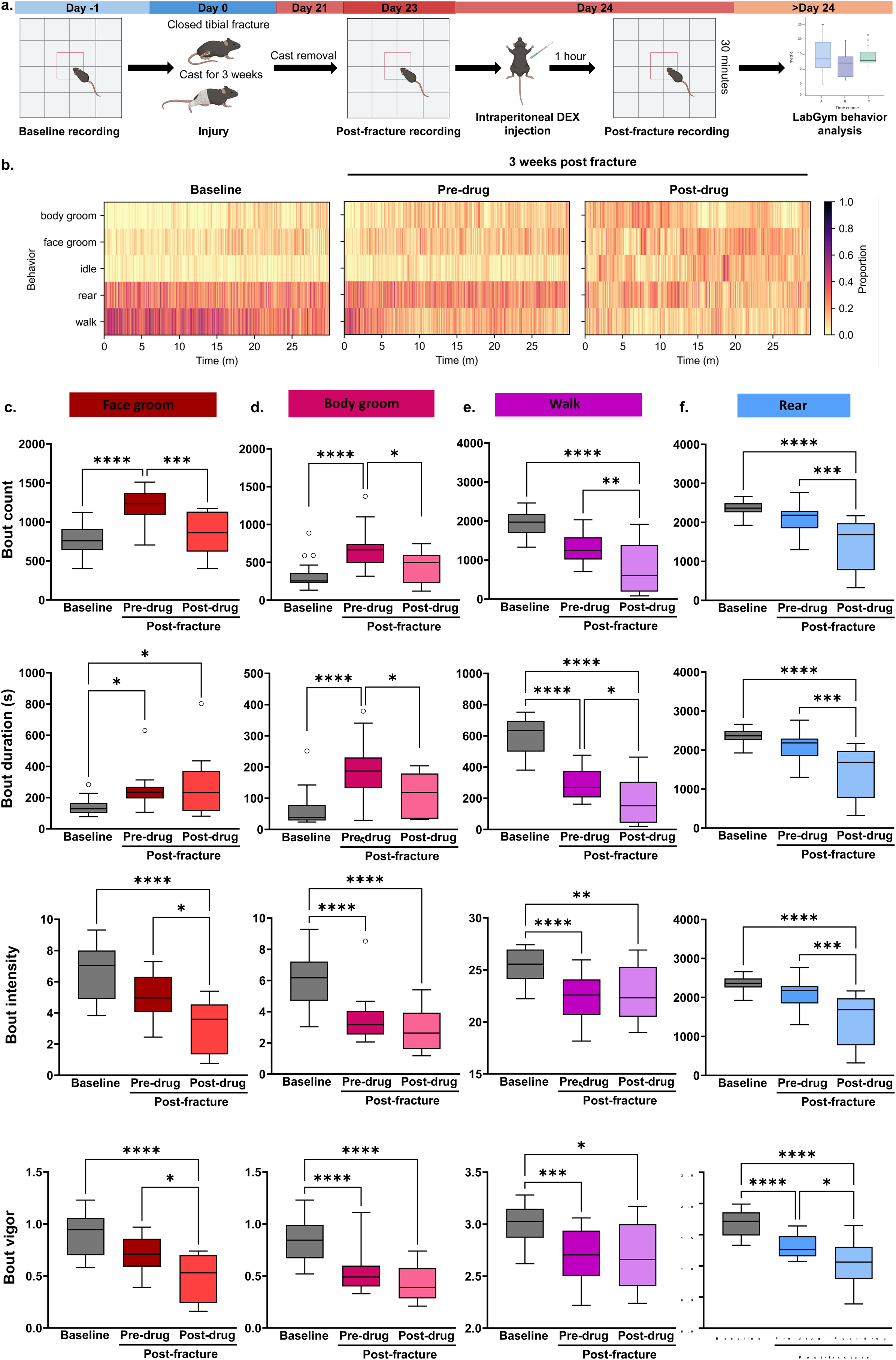
Systemic α2-AR agonism affects naturalistic behaviors in an open field after injury. **a.** Schematic diagram showing experimental timeline for behavior testing and analysis (n= 10 male mice). **b.** Representative raster plots showing proportions of mice performing each behavior at each time point over the course of the analysis. **c.** Intraperitoneal DEX reduces face grooming counts, intensity and vigor, but does not affect face grooming duration in injured mice. **d.** Intraperitoneal DEX reduces body grooming count and duration but does not affect body grooming intensity and vigor in injured mice. **e.** Intraperitoneal DEX further reduces walking frequency and duration but does not affect post injury walking intensity and vigor. **f.** Intraperitoneal DEX further reduces rearing count, duration, intensity and vigor. All behavior data in this figure was analyzed using an ordinary one-way ANOVA with Tukey’s correction (*p<0.05, **p<0.01, ***p<0.001, ****p<0.0001). Data are expressed as the mean ± SEM.

These findings further suggest that DEX treatment at the administered dose may have mildly sedative effects that in turn affect the quality and quantity of naturalistic behaviors in an open field.

### 3.6. α2-AR agonism differentially affects paw licking and grooming in a noxious heat environment after tibial fracture-cast injury

Finally, we conducted an in-depth analysis of α2-AR-mediated effects on injury-induced changes in nocifensive and naturalistic behaviors in a noxious heat environment (Figure 7a). Systemic DEX injection reduced the proportion of injured mice grooming over the course of analysis (Figure 7b). We also observed that systemic DEX administration reversed injury-induced changes in grooming frequency and duration but had no effect on grooming latency (Figure 7c, d, e respectively). Grooming intensity and vigor also decreased after systemic DEX injection (Figure 7f and g respectively). There were no further changes observed in other miscellaneous behaviors, rearing and paw withdrawal after DEX administration (Supplementary figure 7a, b, and c respectively). Although the increase in paw licking latency was reversed by DEX (Figure 7h), no changes were observed in paw licking count, duration, intensity and vigor after injury (Supplementary figure 7d).

**Figure 7.**
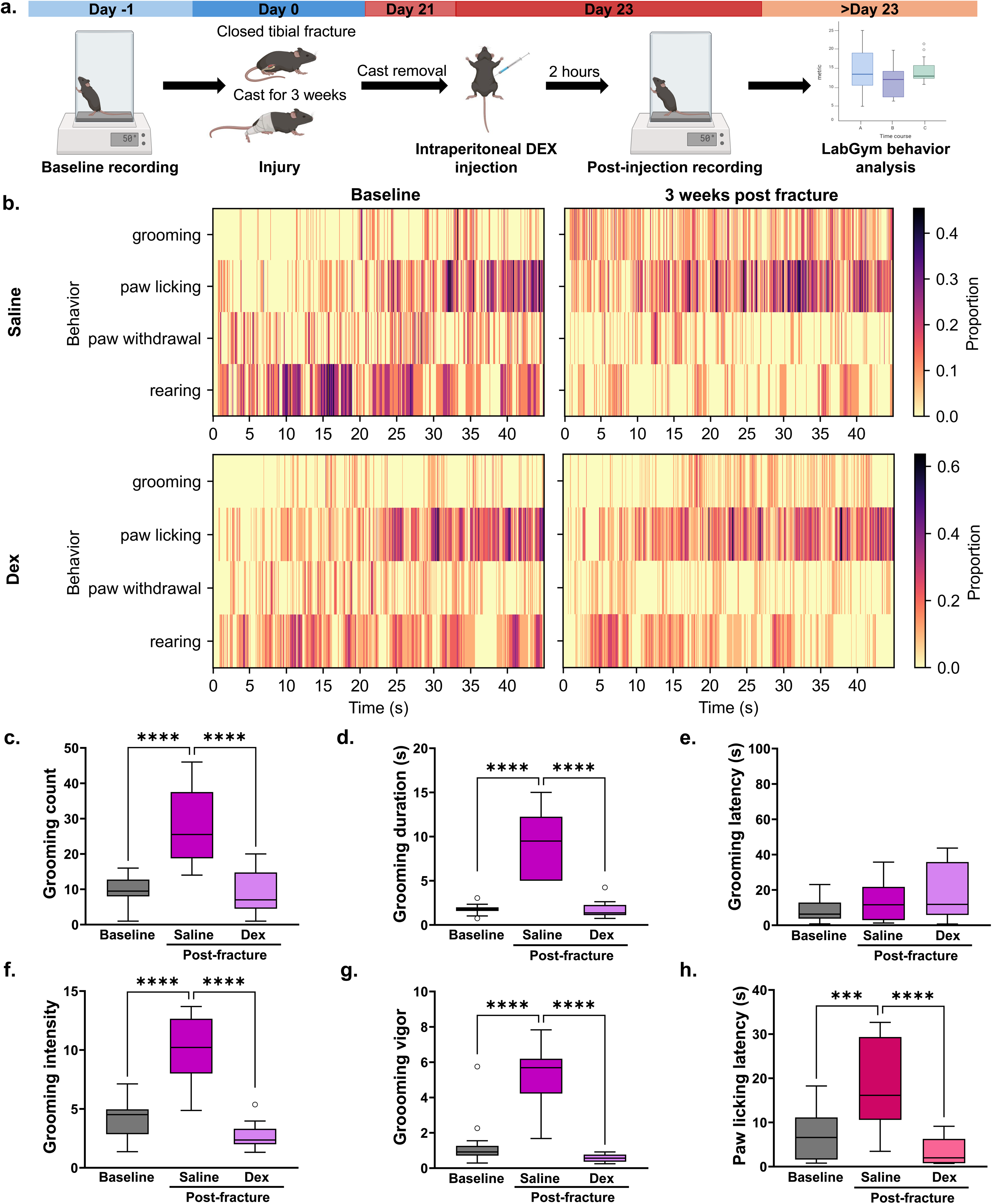
Systemic α2-AR agonism attenuates heat-evoked grooming and paw licking after tibial fracture-cast injury. **a.** Schematic diagram showing experimental timeline for measuring heat evoked pain responses following DEX treatment in injured mice (n= 10 male mice). **b.** Representative raster plots showing proportions of mice performing each behavior at each time point over the course of the analysis. Intraperitoneal DEX reduces grooming count (**c**), duration (**d**), intensity (**f**) and vigor (**g**) but does not affect grooming latency (**e**) in injured mice. **h.** Intraperitoneal DEX restores baseline paw licking latency in injured mice. All behavior data in this figure was analyzed using an ordinary two-way ANOVA with Tukey’s correction (*p<0.05, **p<0.01, ***p<0.001, ****p<0.0001). Data are expressed as the mean ± SEM.

To further delineate the site of action and the relative contributions of central and peripheral α2-AR to the observed effects on pain-related behaviors in a noxious heat environment, we administered α2-AR and α2A-AR antagonists as done previously (Figure 8a). We demonstrate that co-administration of DEX and yohimbine did not result in a reversal of DEX induced decrease in grooming counts, duration, intensity and vigor. Instead, grooming frequency, duration, intensity and vigor were still reduced from baseline in the presence of yohimbine (Figure 8b). Similar results were observed for grooming in a noxious heat environment when DEX was injected in the presence of BRL-44408 maleate. Grooming count, duration, intensity and vigor were all reduced in the presence of BRL-44408 maleate in injured mice (Figure 8c). Injecting DEX in the presence of yohimbine also reduced paw licking counts, duration, latency, vigor and intensity (Figure 8d). Lastly, injecting DEX in the presence of MK467 increased paw withdrawal latency in injured mice (Figure 8e). DEX effects on grooming and paw licking were also preserved when the drug was administered in the presence of MK467 (data not shown).

**Figure 8.**
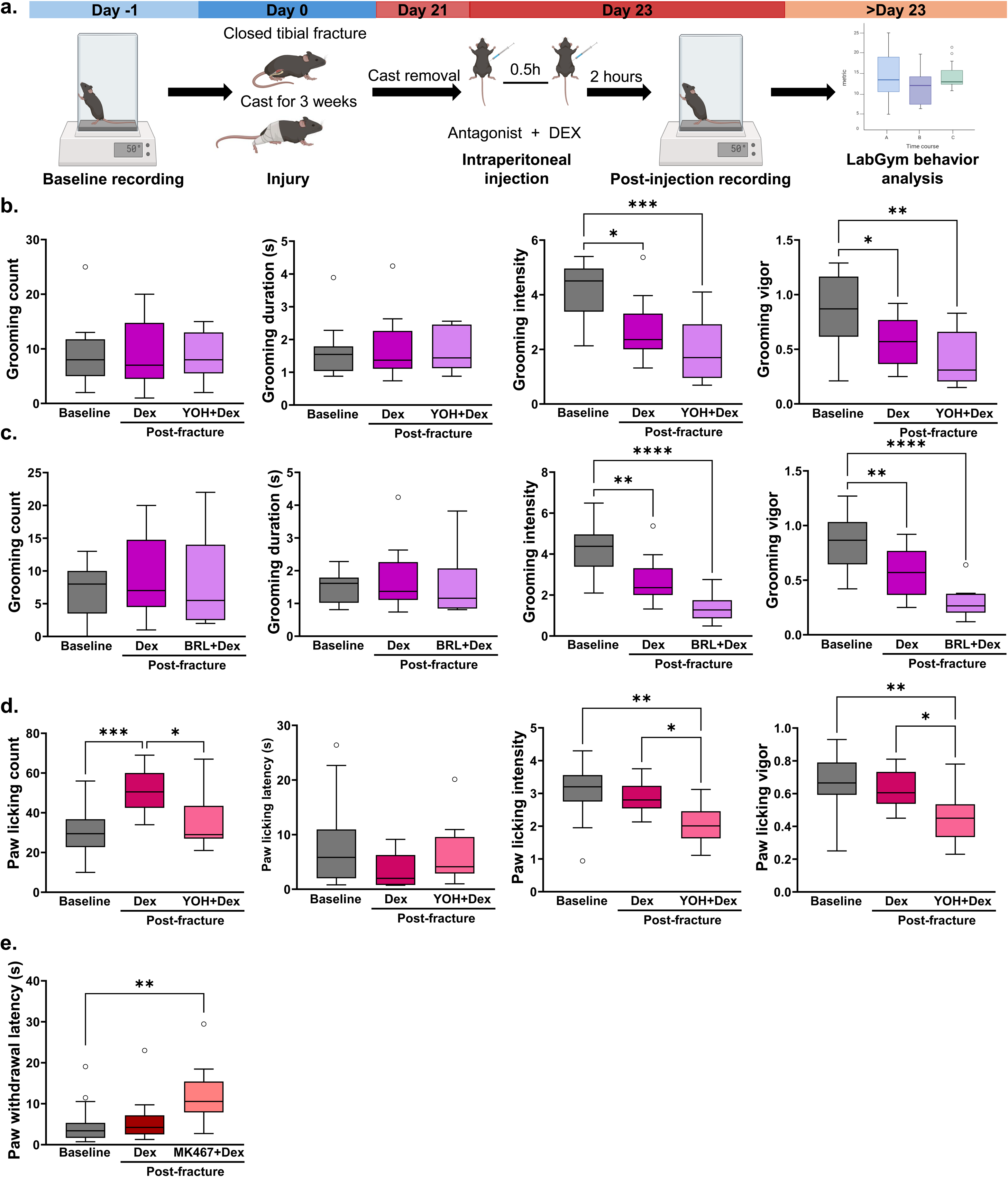
Systemic α2-AR antagonism enhances DEX-induced attenuation of nocifensive and naturalistic behaviors after injury. **a.** Schematic diagram showing experimental timeline for measuring heat evoked pain responses following DEX and antagonist treatment in injured mice. **b.** Administering yohimbine (YOH) with DEX restores baseline grooming frequency and duration but does not affect grooming intensity and vigor in injured mice. **c.** Administering BRL44408 maleate (BRL) with DEX restores baseline grooming frequency and duration but does not affect grooming intensity and vigor in injured mice. **d.** Administering YOH with DEX restores baseline paw licking latency, and reduces paw licking count, intensity and vigor in injured mice. **e.** Administering MK467 with DEX increases paw withdrawal latency in injured mice. All behavior data in this figure was analyzed using an ordinary one-way ANOVA with Tukey’s correction (*p<0.05, **p<0.01, ***p<0.001, ****p<0.0001). Data are expressed as the mean ± SEM.

These findings show that systemic α2A-AR antagonism does not affect DEX-mediated changes in grooming and paw licking, and peripheral α2A-AR antagonism amplifies DEX effects on paw withdrawal in a noxious environment.

## 4. DISCUSSION

### 4.1 A unique ethological signature in the tibial fracture-cast model of complex regional pain syndrome

Chronic post-injury pain has peripheral sensory and central affective components[12,35,71]. Post-injury pain, including CRPS, alters activity in brain regions involved in pain processing such as the anterior cingulate cortex and the amygdala[66]. This in turn can alter stereotypical behaviors like grooming, locomotor and exploratory behavior, and nesting[59]. The ability of pre-clinical models to predict success of novel analgesics is limited[9]. Some of the proposed reasons include evaluation of nociceptive reflex pathways rather than full circuit responses possibly more reflective of “pain.” Automated behavior analysis with tools like MoSeq[68], ARBEL[2], LUPE[30], provides better precision for detecting subtle behavioral changes stemming from full circuit responses in pain models. However, there are significant barriers to integrating automated behavior analysis tools into routine behavior analysis pipelines in pain research labs. Tools like MoSeq[68], ARBEL[2], and LUPE[30] require considerable computational expertise, upfront investment in hardware like high-speed cameras, training personnel, and construction of specialized testing apparatuses. We therefore sought a more user-friendly option for holistic ethological assessment of injured mice. LabGym requires no special equipment, is broadly applicable across behavioral paradigms and requires little to no coding skills[27], making it easier to deploy in labs without extensive computational resources.

Injury-induced changes in stereotypical behaviors may be reflected in their duration, frequency, and intensity. This phenomenon is clearly demonstrated in this study which, for the first time, employs machine learning to illustrate injury-induced changes in naturalistic behaviors in neutral and noxious heat environments in the mouse tibial fracture-cast model of complex regional pain syndrome. It is important to note that the nature of changes in naturalistic behaviors like grooming and rearing can be influenced by the nature of the injury and the environment in which behavior is measured. For example, some researchers report reduced grooming in oxaliplatin-induced neuropathic pain[17], and others report increased grooming in mouse models of inflammatory pain[39] and trigeminal pain[57]. We report an increase in grooming in both neutral and noxious heat environments. Similarly, researchers have found no effect on rearing and exploratory behavior in inflammatory pain induced by Complete Freund’s adjuvant[7], and decreased rearing and exploratory behavior has been observed in mice after fracture[51] that could be due to limited limb function, stress-induced fatigue, and apprehension caused by pain. In our study, mice exhibit reduced rearing and locomotion in an open field after tibial fracture-cast injury. Similar findings have been shown in other mouse fracture models in which the fracture impairs limb function and weight bearing, leading to reduced rearing and walking[43,51]. Mice in our study exhibited reduced rearing and paw withdrawal even when exposed to noxious heat which may be attributed to an aversion for placing weight on the injured limb and vigorous activity. This can be likened to patients with CRPS who often avoid use of their injured limb[25,60]. Perhaps as a coping strategy, in lieu of paw withdrawal, mice licked their paws more frequently and for longer periods of time when placed on a hot plate. We also observed more frequent, intense and vigorous grooming in injured mice when they are placed on a hotplate, which possibly points to heightened stress feeding into an already destabilized neural circuit.

This study provides a comprehensive evaluation of evoked and spontaneous pain in a mouse tibial fracture-cast injury model of CRPS. It combines previously documented changes in reflexive pain and autonomic signs with a detailed analysis of the changes in stereotypical mouse behaviors in neutral and noxious heat environments after injury.

### 4.2 Peripheral vs. central action of DEX after tibial fracture-cast injury

We used DEX to characterize the effect of targeting α2-ARs on evoked and spontaneous pain behaviors, motor function and anxiety-like behaviors in injured mice. By reducing the release of norepinephrine, DEX dampens neuronal excitability in pain pathways, resulting in analgesia, anxiolysis, and, at higher doses, sedation[74]. As in prior studies in other models of pain[15,32,38,40,74,75], DEX reduced mechanical allodynia in the tibial fracture-cast model of CRPS. DEX also reduced paw temperature after injury. Both temperature increase and decrease have been associated with DEX in humans and the underlying mechanisms in either case are still unclear[11]. In rodents, DEX reduces body temperature primarily through modulating thermoregulation in the hypothalamus[8,41,47]. However, peripheral vascular α2-ARs have also been implicated in DEX-mediated hypothermia[1,20] and could contribute to our finding. DEX also has anti-inflammatory properties that could indirectly result in paw temperature reduction[73]. The analgesic and anxiolytic effects of DEX may also normalize naturalistic behaviors that are altered by chronic post-injury pain. Holistic assessment with LabGym allowed us to quantify these changes in fractured mice. We observed improved grooming in injured mice in both neutral and noxious heat environments. We did not, however, see an improvement in locomotion or rearing. Instead, locomotion and rearing were further reduced after DEX treatment. This observation points to sedation which could mask any alleviating effect on locomotion and rearing.

We next interrogated the relative contribution of peripheral and central α2-ARs to DEX action on mechanical allodynia and naturalistic behaviors in a noxious heat environment. We hypothesized that the modulating effects of DEX in our study resulted from combined activation of central and peripheral α2-ARs and used selective antagonists to test this hypothesis. We discovered that mechanical allodynia after tibial fracture-cast injury is primarily mediated by central α2A-ARs. Conversely, systemic and peripheral antagonism of α2A-ARs did not mitigate DEX-mediated effects on naturalistic behaviors in a noxious heat environment. This could indicate that reflexive pain after tibial fracture-cast injury is not mediated by true sympathetic nervous system responses but rather only by somatic α2-AR. Additionally, persistent DEX effects on naturalistic behaviors with concomitant α2A-AR antagonism in our study open the door to several possibilities regarding mechanisms of DEX analgesia after tibial fracture-cast injury. Emerging evidence suggests that in addition to α2A-AR activation, DEX may engage additional mediators and modulatory pathways to achieve analgesia. For example, DEX has been reported to interact with imidazoline receptors that are thought to participate in central sympathetic modulation and might help dampen the stress response after injury[24,31,61]. Studies have also shown that DEX acts as an anti-inflammatory by reducing the release of pro-inflammatory cytokines and modulating neuroinflammatory responses [37,69,73], a response that may be an indirect effect of DEX action on sympathetic tone. Likewise, DEX has been shown to exert a neuroprotective role by modulating glial activity[52], which could also contribute to DEX analgesia. Lastly, DEX might influence other neurotransmitter systems such as opioid systems[5,19,34], which further complicates the delineation of a precise mechanism for DEX analgesia. This is in no way an exhaustive list and further investigation is warranted to provide deeper insight into DEX’s mechanism of action which in turn would allow optimization of DEX’s therapeutic benefits.

This unique ethological signature which incorporates aspects of behavior that are reflective of the human condition, provides a platform for development for mechanism-based treatment, as shown by our proof-of-concept using DEX.

## 5. STUDY LIMITATIONS

First, experiments in this study were conducted only in male mice. We have previously noted similar timelines to injury resolution and involvement of microglia in both male and female mice in this model. It is therefore possible that the role of α2-AR is also conserved across sexes. Secondly, mice were tested at a single timepoint after tibial fracture-cast surgery which was selected given the known acute-to-chronic or peripheral-to-central transition at that timepoint. Performing such analyses at multiple timepoints after injury would provide a more holistic view of pain behaviors in CRPS and the viability of α2-AR-targeting for chronic pain treatment. Finally, drug experiments involved a single drug treatment followed by behavior assessments. Continuous infusions or repeated dosing paradigms may have different effects on pain behaviors in this mouse model.

## AUTHORSHIP CONTRIBUTION STATEMENT

Gabriella P. B. Muwanga: Writing – review & editing, Writing – original draft, Figure generation, Experimental design, Data collection and analysis, Funding acquisition, Conceptualization.

Amanda Pang: Writing – review & editing, Data collection and analysis.

Sedona N. Ewbank: Writing – review & editing, Data analysis.

Janelle Siliezar-Doyle: Writing – review & editing, Data collection.

Amy R. Nippert: Writing – review & editing, Data collection.

Raag D. Airan: Writing – review & editing, Conceptualization

Vivianne L. Tawfik: Writing – review & editing, Project administration, Funding acquisition, Conceptualization.

## DECLARATION OF GENERATIVE AI AND AI-ASSISTED TECHNOLOGIES IN THE WRITING PROCESS

The authors used ChatGPT 4.0 for proofreading parts of the introduction and discussion section during the preparation of this work. After using this tool, we reviewed and edited the content as needed and take full responsibility for the content of the publication.

## ACKNOWLEDGEMENTS

We sincerely thank Kanchan Sinha Roy for his contributions to initial experiments and Lauren Donovan for technical assistance in earlier iterations of this project. We also extend our gratitude to the entire Tawfik and Airan labs for invaluable feedback and discussion.

## DATA AVAILABILITY

Data generated in this study will be uploaded to a public drive for easy access.

