## Supplemental Figures for "Ethological profiling of pain and analgesia in a mouse model of complex regional pain syndrome"

**Supplementary figure 1. LabGym open field analysis model achieves high precision, recall, and F1 scores for all behaviors of interest. a.** Open field analysis model identifies idling with high precision, recall, and F1 scores in uninjured (naïve) and injured mice (Uninjured: precision = 1.0, recall = 0.89, F1 score = 0.94; injured: precision = 0.95, recall = 0.92, F1 score = 0.95; mixed: precision = 0.94, recall = 0.91, F1 score = 0.93). **b.** Open field analysis model identifies walking with high precision, recall, and F1 scores in uninjured (naïve) and injured mice (Uninjured: precision = 0.97, recall = 0.99, F1 score = 0.98; injured: precision = 0.97, recall = 0.98, F1 score = 0.98; mixed: precision= 0.98, recall = 0.98, F1 score= 0.98). **c.** Open field analysis model identifies rearing with high precision, recall, and F1 scores in uninjured and injured mice (Uninjured: precision = 0.93, recall = 0.96, F1 score = 0.95; injured: precision = 0.92, recall = 0.9, F1 score = 0.91; mixed: precision = 0.95, recall = 0.93, F1 score = 0.94). **d.** Open field analysis model identifies body grooming with high precision, recall, and F1 scores in naïve and injured mice (Uninjured: precision = 0.98, recall = 0.92, F1 score = 0.95; injured: precision = 0.98, recall = 0.95, F1 score = 0.97; mixed: precision = 0.98, recall = 0.94, F1 score = 0.96). **e.** Open field analysis model identifies face grooming with high precision, recall, and F1 scores in naïve and injured mice (Uninjured: precision = 0.96, recall = 0.98, F1 score = 0.97; injured: precision = 0.83, recall = 0.89, F1 score = 0.86; mixed: precision = 0.86, recall = 0.95, F1 score = 0.9).

**Supplementary figure 2. Tibial fracture-cast injury increases time spent idle but does not affect pain-related behavior latencies in an open field. a.** No statistically significant changes in open field behavior latencies are observed in injured mice three weeks post fracture. **b.** Tibial fracture-injury increases time spent idle but does not affect how often or how long it takes for injured mice to become idle. All behavior data shown in this figure was analyzed using Student's t-test (*p<0.05, **p<0.01, ***p<0.001, ***p<0.0001). Data are expressed as the mean ± SEM.

**Supplementary figure 3. LabGym hotplate analysis model achieves high precision, recall, and F1 scores for all behaviors of interest. a.** Hotplate analysis model identifies miscellaneous behaviors with high precision, recall, and F1 scores in uninjured (naïve) and injured mice (Uninjured: precision = 0.86, recall = 0.92, F1 score = 0.89; injured: precision = 0.97, recall = 0.93, F1 score = 0.95; mixed: precision = 0.91, recall = 0.91, F1 score = 0.92). **b.** Hotplate analysis model identifies grooming with high precision, recall, and F1 scores in uninjured (naïve) and injured mice (Uninjured: precision = 0.87, recall = 0.81, F1 score = 0.84; injured: precision = 0.95, recall = 0.86, F1 score = 0.9; mixed: precision= 0.91, recall = 0.84, F1 score= 0.87). **c.** Hotplate analysis model identifies rearing with high precision, recall, and F1 scores in uninjured and injured mice (Uninjured: precision = 0.95, recall = 0.95, F1 score = 0.95; injured: precision = 0.94, recall = 0.96, F1 score = 0.95; mixed: precision = 0.95, recall = 0.96, F1 score = 0.95). **d.** Hotplate analysis model identifies paw withdrawal with high precision, recall, and F1 scores in naïve and injured mice (Uninjured: precision = 0.92, recall = 0.76, F1 score = 0.83; injured: precision = 0.95, recall = 0.9, F1 score = 0.92; mixed: precision = 0.93, recall = 0.8, F1 score = 0.86). **e.** Hotplate analysis model identifies paw licking with high precision, recall, and F1 scores in naïve and injured mice (Uninjured: precision = 0.88, recall = 0.93, F1 score = 0.9; injured: precision = 0.72, recall = 0.95, F1 score = 0.82; mixed: precision = 0.82, recall = 0.93, F1 score = 0.87).

**Supplementary figure 4. Tibial fracture-cast injury in mice does not significantly affect every heat induced behavior metric recorded on a hot plate. a.** No statistically significant changes in thermal induced paw withdrawal, paw licking, grooming, and jumping latencies are observed in CRPS mice three weeks post fracture. **b.** No statistically significant changes in thermal induced paw withdrawal, paw licking, grooming, and rearing frequency are observed in CRPS mice three weeks post fracture. **c.** No statistically significant changes in thermal induced paw licking and jumping intensity are observed in CRPS mice three weeks post fracture. **d.** No statistically significant changes in thermal induced paw licking and jumping vigor are observed in CRPS mice three weeks post fracture. **e.** No statistically significant changes in thermal induced paw withdrawal, paw licking, and rearing duration are observed in CRPS mice three weeks post fracture. All behavior data shown in this figure was analyzed using Student's t-test (*p<0.05, **p<0.01, ***p<0.001, ***p<0.0001). Data are expressed as the mean ± SEM.

**Supplemental figure 5. DEX reduces motion in an open field after tibial fracture-cast injury. a.** Schematic diagram showing experimental timeline for open field behavior testing and analysis (n= 10 male mice per group). **b.** Representative path plots showing post injury and post saline injection changes in motion in injured mice. **c.** Representative path plots showing post injury and post DEX injection changes in motion in injured mice. **d.** Intraperitoneal DEX does not significantly reduce the distance traveled in an open field in injured mice. **e.** Intraperitoneal DEX does not significantly reduce average speed traveled in an open field in injured mice. **f.** Intraperitoneal DEX does not affect time spent in the center zone after injury. **g.** Intraperitoneal DEX does not affect time spent in the thigmotaxis zone after injury. **h.** Intraperitoneal DEX increases the total time spent immobile by injured mice in an open field. All behavior data in this figure was analyzed using an ordinary two-way ANOVA with Tukey's correction (*p<0.05, **p<0.01, ***p<0.001, ****p<0.0001). Data are expressed as the mean ± SEM.

**Supplemental figure 6. DEX increases walking latency but does not affect other pain-related behavior latencies and idling behavior dynamics in an open field after injury. a.** DEX increases walking latency but does not affect grooming or rearing latencies after injury. **b.** DEX does not affect idle bout counts, duration after injury. All behavior data shown in this figure was analyzed using an ordinary one-way ANOVA with Tukey's correction (*p<0.05, **p<0.01, ***p<0.001, ****p<0.0001). Data are expressed as the mean ± SEM.

**Supplemental figure 7. DEX does not affect rearing, paw withdrawal, or other miscellaneous behaviors in a noxious heat environment after injury. a.** DEX does not affect thermal-induced changes in miscellaneous behaviors after tibial fracture-cast injury. **b.** DEX does not affect thermal-induced changes in rearing after tibial fracture-cast injury. **c.** DEX does not affect thermal-induced changes in paw withdrawal after tibial fracture-cast injury. **d.** DEX does not affect thermal induced paw licking and jumping intensity in CRPS mice three weeks post fracture. **e.** DEX does not affect thermal-induced changes paw licking count, duration, intensity and vigor. All behavior data shown in this figure was analyzed using an ordinary one-way ANOVA with Tukey's correction (*p<0.05, **p<0.01, ***p<0.001, ****p<0.0001). Data are expressed as the mean ± SEM.

SUPPLEMENTAL FIGURE 1

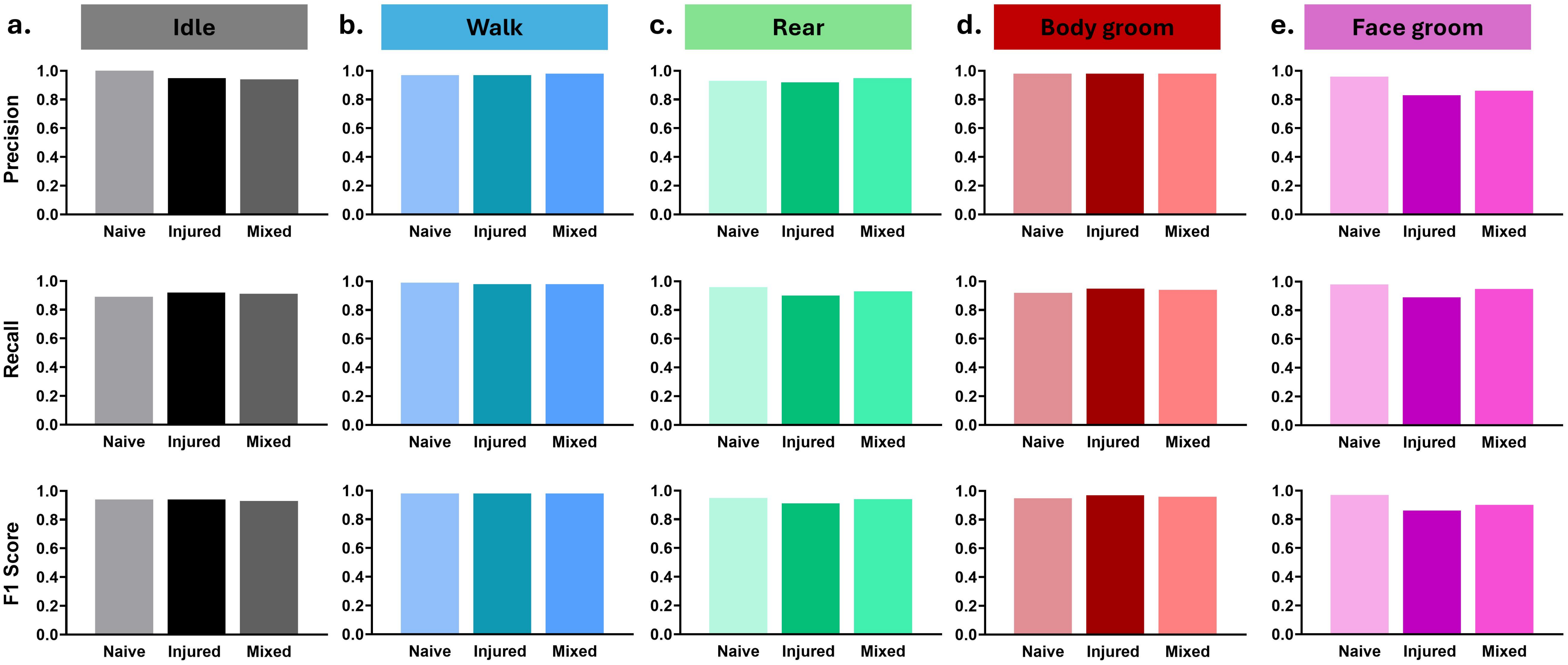

SUPPLEMENTAL FIGURE 2

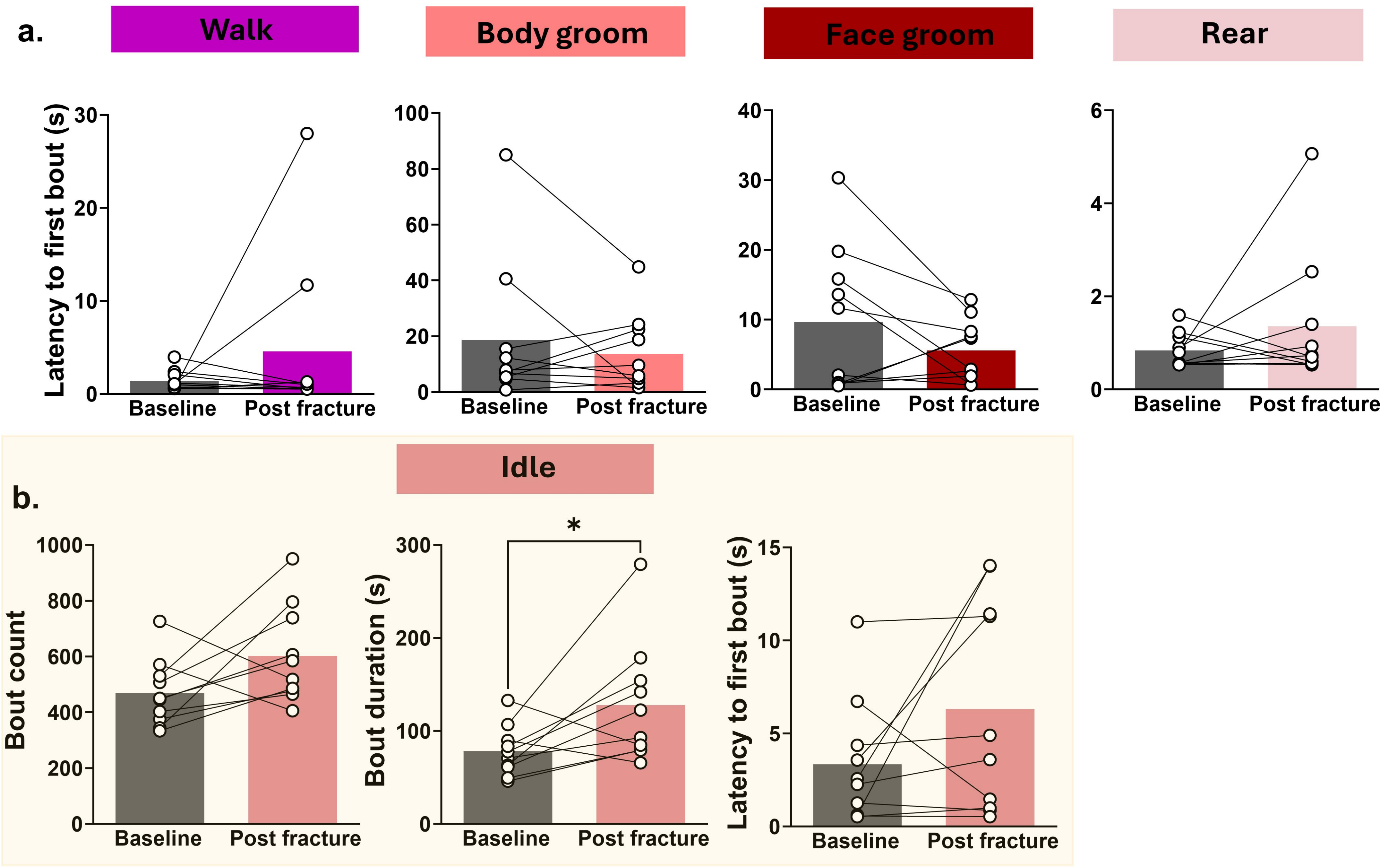

SUPPLEMENTAL FIGURE 3

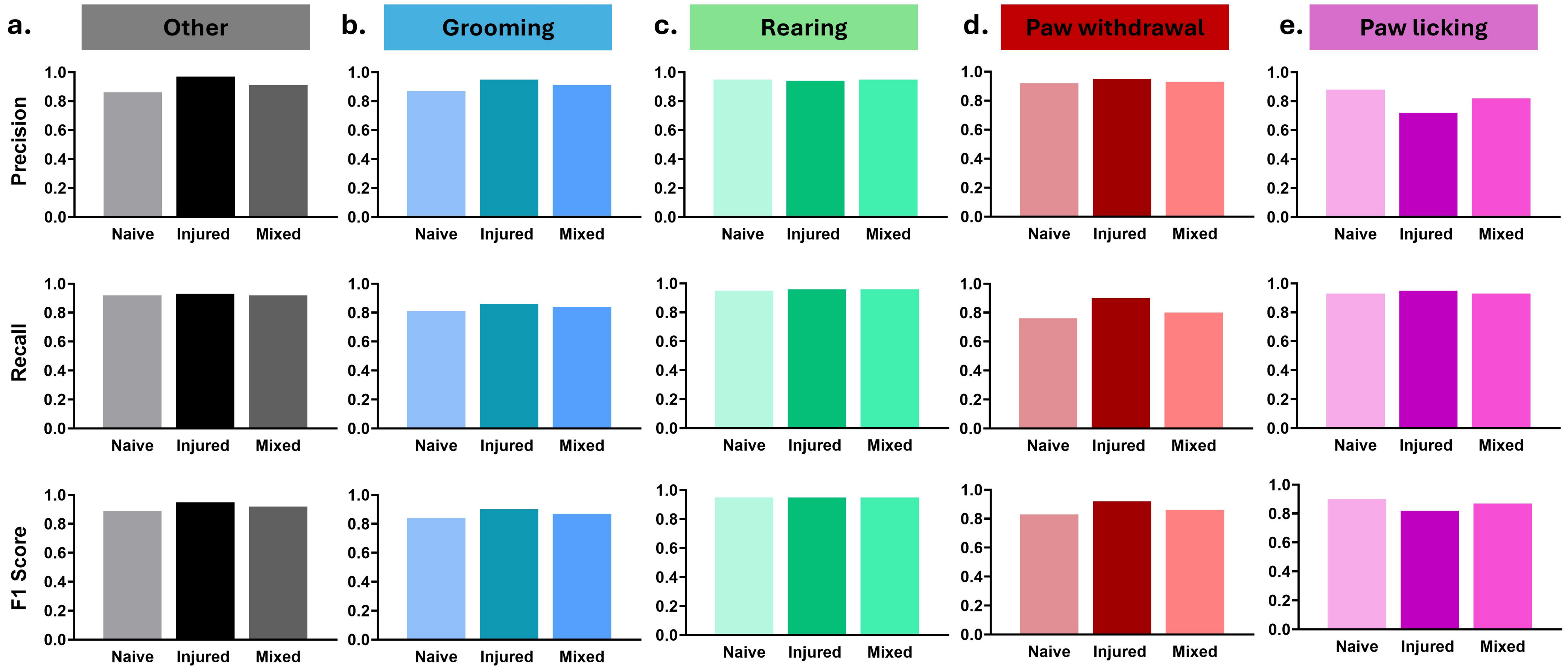

SUPPLEMENTAL FIGURE 4

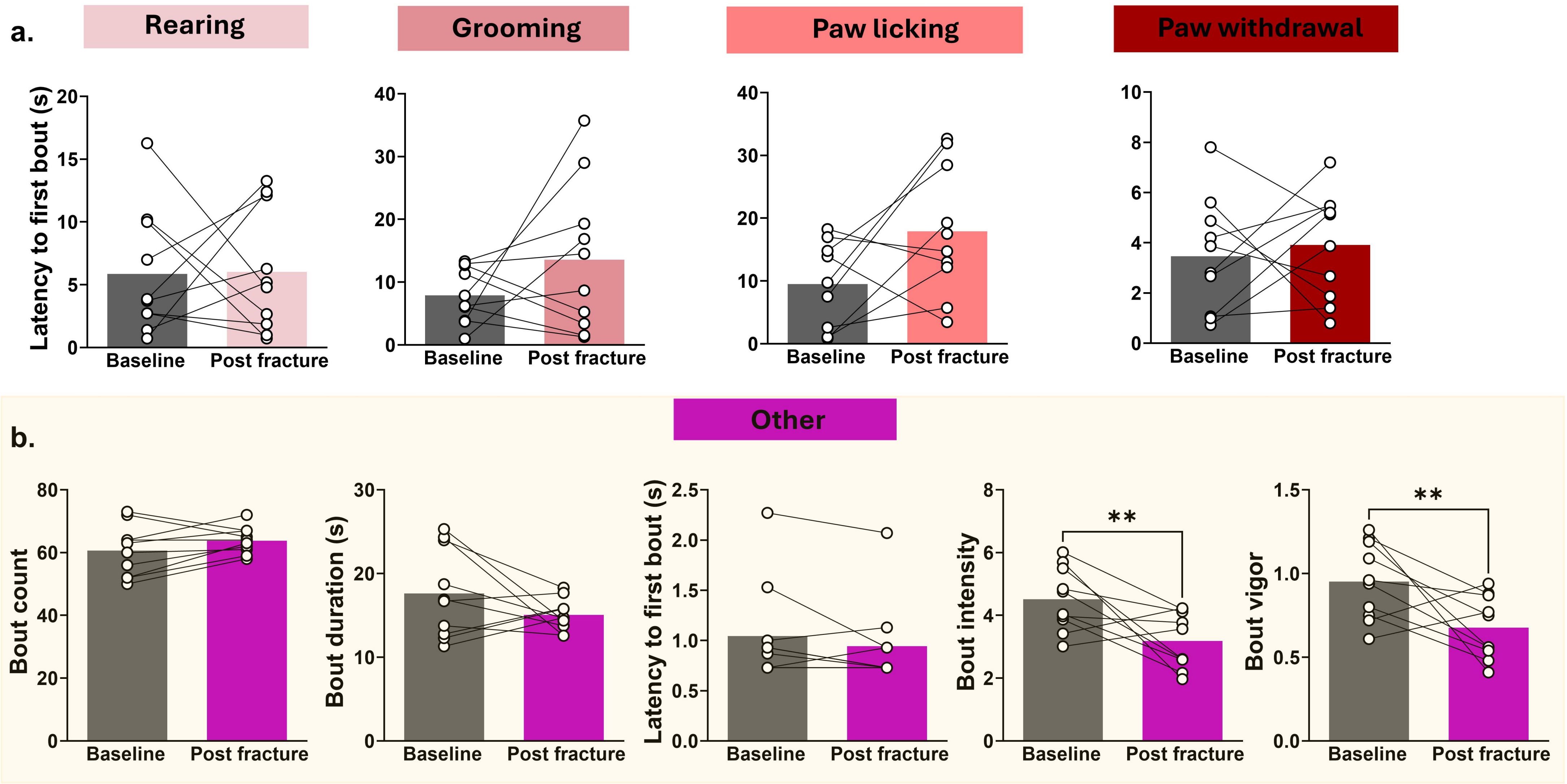

### SUPPLEMENTAL FIGURE 5

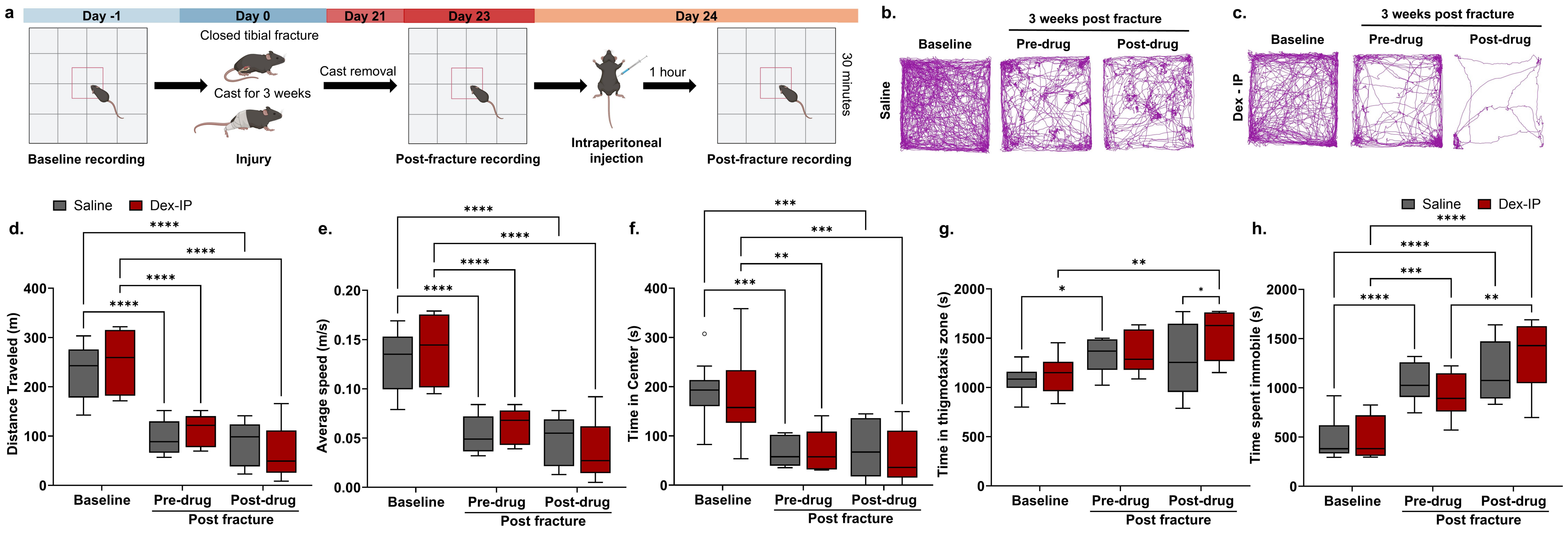

SUPPLEMENTAL FIGURE 6

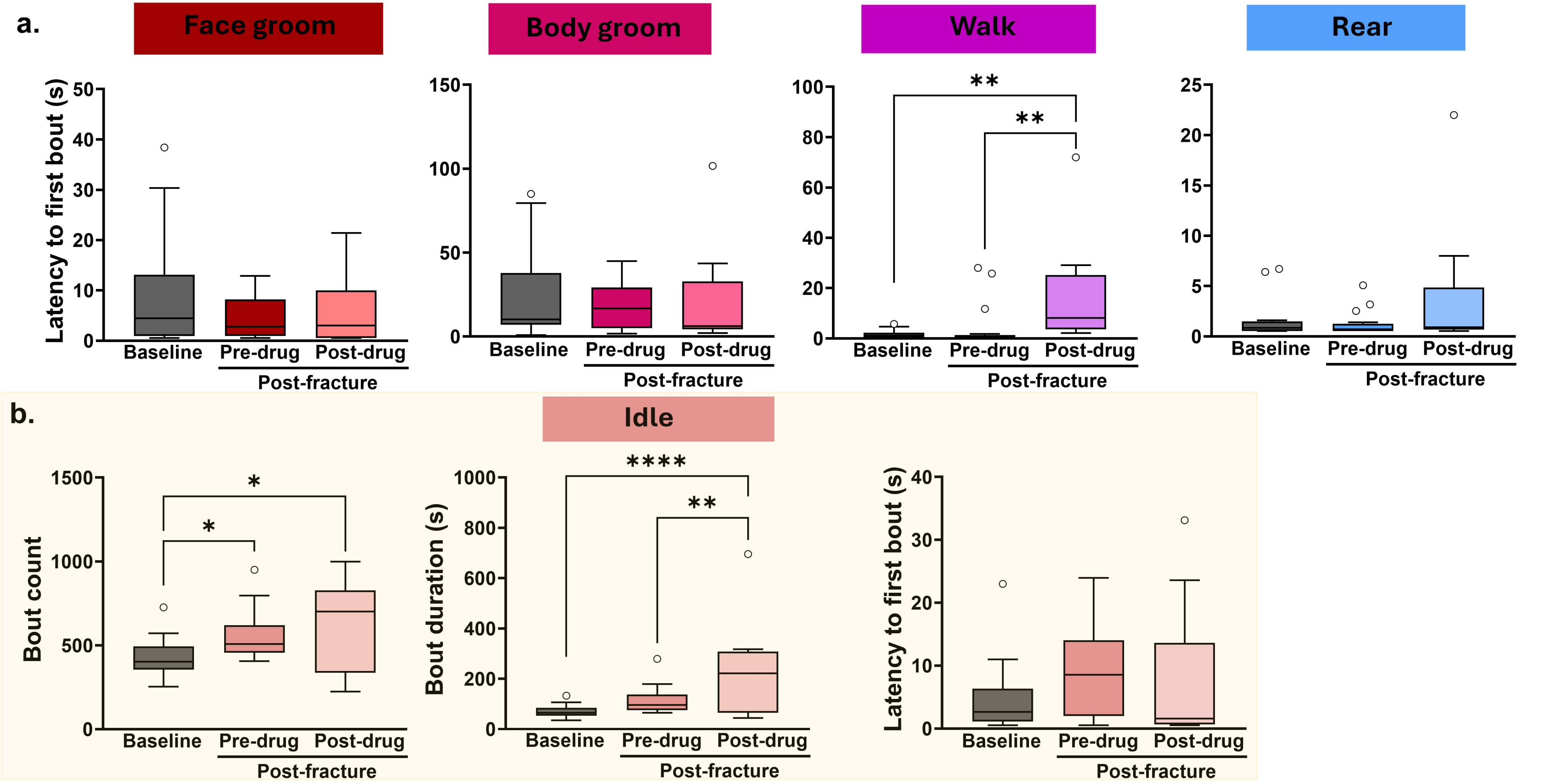

#### SUPPLEMENTAL FIGURE 7

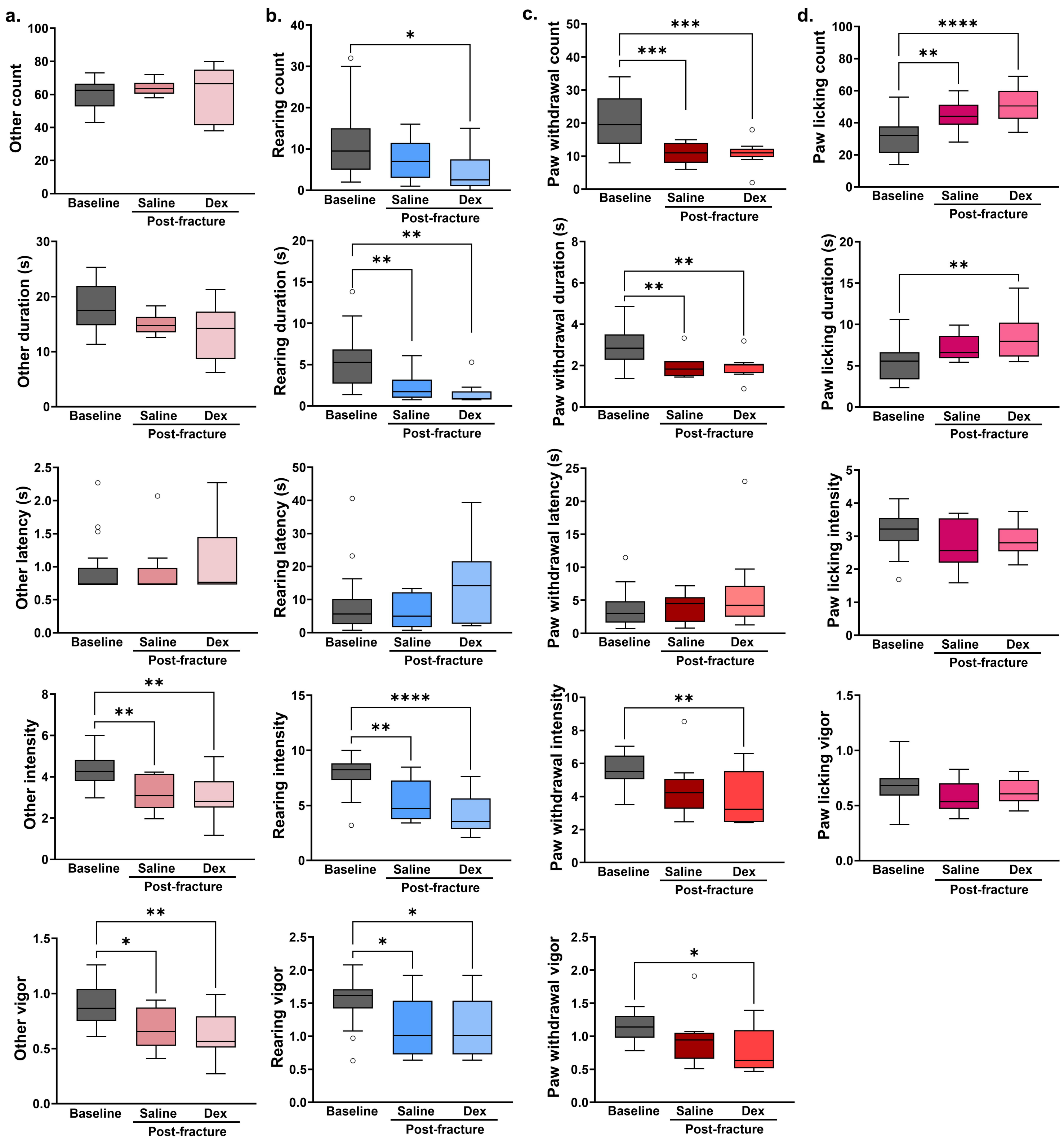
